# Hebbian Learning in a Random Network Captures Selectivity Properties of Prefrontal Cortex

**DOI:** 10.1101/133025

**Authors:** Grace W. Lindsay, Mattia Rigotti, Melissa R. Warden, Earl K. Miller, Stefano Fusi

## Abstract

Complex cognitive behaviors, such as context-switching and rule-following, are thought to be supported by prefrontal cortex (PFC). Neural activity in PFC must thus be specialized to specific tasks while retaining flexibility. Nonlinear ‘mixed’ selectivity is an important neurophysiological trait for enabling complex and context-dependent behaviors. Here we investigate (1) the extent to which PFC exhibits computationally relevant properties such as mixed selectivity and (2) how such properties could arise via circuit mechanisms. We show that PFC cells recorded from male and female rhesus macaques during a complex task show a moderate level of specialization and structure that is not replicated by a model wherein cells receive random feedforward inputs. While random connectivity can be effective at generating mixed selectivity, the data shows significantly more mixed selectivity than predicted by a model with otherwise matched parameters. A simple Hebbian learning rule applied to the random connectivity, however, increases mixed selectivity and allows the model to match the data more accurately. To explain how learning achieves this, we provide analysis along with a clear geometric interpretation of the impact of learning on selectivity. After learning, the model also matches the data on measures of noise, response density, clustering, and the distribution of selectivities. Of two styles of Hebbian learning tested, the simpler and more biologically plausible option better matches the data. These modeling results give intuition about how neural properties important for cognition can arise in a circuit and make clear experimental predictions regarding how various measures of selectivity would evolve during animal training.

**Significance Statement:** Prefrontal cortex (PFC) is a brain region believed to support the ability of animals to engage in complex behavior. How neurons in this area respond to stimuli—and in particular, to combinations of stimuli (”mixed selectivity”)—is a topic of interest. Despite the fact that models with random feedforward connectivity are capable of creating computationally-relevant mixed selectivity, such a model does not match the levels of mixed selectivity seen in the data analyzed in this study. Adding simple Hebbian learning to the model increases mixed selectivity to the correct level and makes the model match the data on several other relevant measures. This study thus offers predictions on how mixed selectivity and other properties evolve with training.

## 1. Introduction

The ability to execute complex, context-dependent behavior is evolutionarily valuable and ethologically observed (Rendall et al., 1999; Kalin et al., 1991). How the brain carries out complex behaviors is thus the topic of many neuroscientific studies. A region of focus is the prefrontal cortex (PFC), (Botvinick, 2008; Waskom et al., 2014; Miller and Cohen, 2001; Duncan, 2001), as lesion (Szczepanski and Knight, 2014) and imaging (Miller and D’Esposito, 2005; Bugatus et al., 2017) studies have implied its role in complex cognitive tasks. As a result, several theories have been put forth to explain how PFC can support complexity on the computational and neural levels (Miller and Cohen, 2001; Wood and Grafman, 2003; Fusi et al., 2016).

Observing the selectivity profiles of its constituent cells is a common way to investigate a neural population’s role in a computation. In its simplest form, this involves modeling a neuron’s firing rate as a function of a single stimulus, or, perhaps, an additive function of multiple stimuli (Sahani and Linden, 2003; Duhamel et al., 1998; Moser et al., 2008). More recently, however, the role of neurons that combine inputs in a nonlinear way has been investigated (Rigotti et al., 2013; Mante et al., 2013; Stokes et al., 2013; Pagan et al., 2013; Meister et al., 2013; Raposo et al., 2014; Fusi et al., 2016), often in PFC. Rather than responding only to changes in one input, or to changes in multiple inputs in a linear way, neurons with nonlinear mixed selectivity have firing rate responses that are a nonlinear function of two or more inputs (Figure 1B). Cells with this selectivity (which we call simply “mixed”) are important for population coding because of their effect on the dimensionality of the representation: they increase the dimensionality of the population response, which increases the number of patterns that a linear classifier can read out. This means that arbitrary combinations of inputs can be mapped to arbitrary outputs. In relation to complex behaviors, mixed selectivity allows for a change in context, for example, to lead to different behavioral outputs, even if stimulus inputs are the same. For more on the benefits of mixed selectivity, see Fusi et al. (2016).

**Figure 1:**
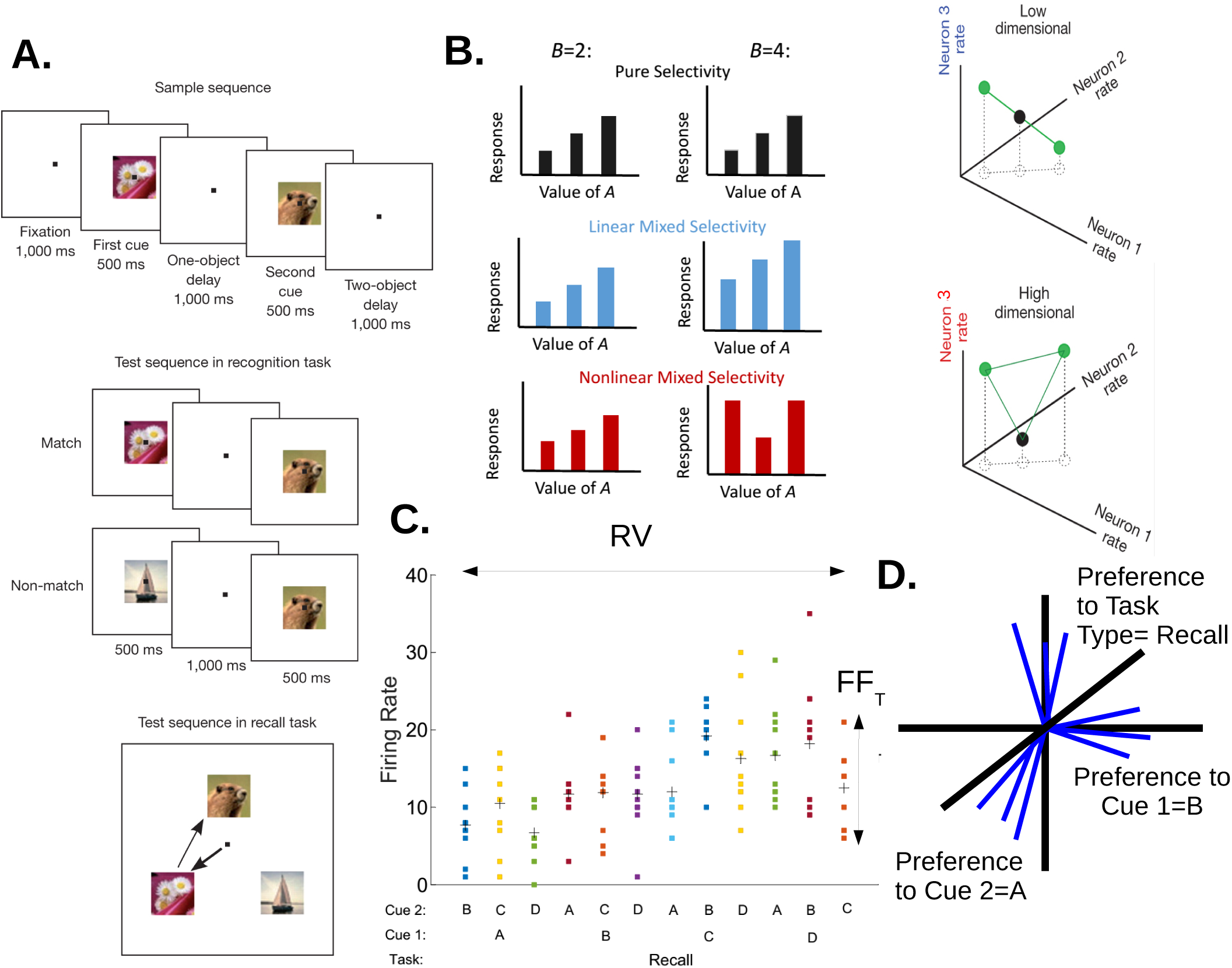
Description of prefrontal cortex data and relevant measures of selectivity A.) Task Design. In both task types, the animal fixated as two image cues were shown in sequence. After a delay the animal had to either indicate that a second presented sequence matched the first or not (”recognition”) or saccade to the two images in correct order from a selection of three images (”recall”). B.) What nonlinear mixed selectivity can look like in neural responses and its impact on computation. The bar graphs on the left depict three different imagined neurons and their responses to combinations of two task variables A and B. The black neuron has selectivity only to A, as its responses are invariant to changes in B. The blue neuron has linear mixed selectivity to A and B: its responses to different values of A are affected by the value of B, but in a purely additive way. The red neuron has nonlinear mixed selectivity: its responses to A are impacted nonlinearly by a change in the value of B. The figures on the right show how including a cell with nonlinear mixed selectivity in a population increases the dimensionality of the representation. With the nonlinearly-selective cell (bottom), the black dot can be separated with a line from the green dots. Without it (top), it cannot. C.) A depiction of measures of trial-to-trial noise (*FF*_*T*_) and the distribution of responses across conditions (RV). The x-axis labels the condition, each dot is the firing rate for an individual trial and the crosses are condition means used for calculating RV (data from a real neuron; recognition task not shown). D.) Conceptual depiction of the clustering measure. Each cell was represented as a vector (blue) in a space wherein the axes (black) represent preference for task variable identities, as determined by the coefficients from a GLM (only three are shown here). The clustering measure determines if these vectors are uniformly distributed.

Theoretical work on how these properties can arise on a circuit level shows that random connectivity is surprisingly efficient at increasing the dimensionality of the neural representation (Jaeger and Haas, 2004; Maass et al., 2002; Buonomano and Maass, 2009; Rigotti et al., 2010; Barak et al., 2013; Babadi and Sompolinsky, 2014; Litwin-Kumar et al., 2017). This means that mixed selectivity can be observed even without learning. However, learning can greatly improve the ability of a linear readout to generalize and hence to make the readout response more robust to noise and variations in the sensory inputs (see e.g. Fusi et al. (2016)). The ideal situation would be one in which a neural population represents only the task relevant variables and the representation has the maximal dimensionality. In brain areas like PFC, where there is a huge convergence of inputs from many other brain areas, it might be important to bias the mixed selectivity representations toward the task relevant variables, which can be achieved only with learning.

In this study, we characterize the response of a population of PFC cells in terms of the distribution of linear and nonlinear selectivity, the response density, and the clustering of selectivities. All these properties characterize the dimensionality of neural representations and are important for the readout performance. As described above, nonlinear mixed selectivity is important for increasing dimensionality. High dimensionality, however, also requires a diversity of responses. We studied this by determining how the preference to different stimuli are distributed across the population. In some lower sensory areas, cells tend to be categorizable—that is, there are groups of cells that display similar preference profiles (Goard et al., 2016). More associative areas tend to lose this clustering of cell types. Such categories may be useful when an area is specialized for a given task, but diversity is needed for flexibility (Raposo et al., 2014).

After characterizing the PFC response, we show that a model with random connectivity can only partially explain the PFC representation. However, with a relatively small deviation from random connectivity—obtained with a simple form of Hebbian learning that is characterized by only two parameters—the model describes the data significantly better.

## 2. Materials and Methods

## 2.1 Task Design

The data used in this study comes from previously published work (Warden and Miller, 2010). In brief, two monkeys performed two variants of a delayed match-to-sample task (Figure 1A). In both task types, after initial fixation, two image cues (chosen from four possible) were presented in sequence for 500ms each with a 1000ms delay period in between the first and second cue. After a second delay period also lasting 1000ms, one of two events occurred, depending on the task type. In the recognition task, another sequence of two images was shown and the monkey was instructed to release a bar if this test sequence matched the initial sample sequence. In the recall task, an array of three images appeared on the screen, and the monkey had to saccade to the two images from the sample sequence in the correct order. Blocks of recall and recognition tasks were interleaved during each recording session. Given that each sequence had two different image cues chosen from the four total image identity options and that there were two task types, the total number of conditions was 4 × 3 × 2 = 24.

## 2.2 Neural Data

Recordings were made using grids with 1 mm spacing (Crist Instrument) and custom-made independently moveable microdrives to lower eight dura-puncturing Epoxylite-coated tungsten microelectrodes (FHC) until single neurons were isolated. Cells were recorded from two adult rhesus monkeys (Macaca mulatta), one female and one male, and combined for analysis. No attempt was made to pre-screen neurons, and a total of 248 neurons were recorded (with each neuron observed under both task types).

For the purposes of this study, firing rates for each neuron were calculated as the total number of spikes during the later 900ms of the second delay period, as it was at this point that the identities of all task variables were known. Any cells that did not have at least 10 trials for each condition or did not have a mean firing rate of at least 1 spike/sec as averaged over all trials and conditions were discarded. This left 90 cells.

## 2.3 Fano Factor Measurements

Noise is an important variable when measuring selectivity. High noise levels require stronger tuning signals in order to be useful for downstream areas, and to reach significance in statistical testing. Thus, any model attempting to match the selectivity profile of a population must be constrained to have the same level of noise. Here, we measure noise as the Fano Factor (variance divided by mean) of each cell’s activity across trials for each condition (spike count taken from later 900ms of the two-object delay). This gives 24 values per cell. This is the trial Fano Factor. Averaging over conditions gives one trial Fano Factor value per cell, and averaging over cells gives a single number representing the average noise level of the network. Unless otherwise stated, *FF*_*T*_ refers to this network averaged measure.

Another measure of interest is how a neuron’s response is distributed across conditions. Do neurons respond differentially to a small number of conditions (i.e., a sparse response), or is the distribution more flat? To measure this, the firing rate for each condition (averaged across trials) was calculated for each neuron and the Fano Factor was calculated across conditions. In this case, a large value means that some conditions elicit a very different response than others, while a small value suggests the responses across conditions are more similar. We call this value the response variability, or RV. Averaging across all cells gives the response variability of the network.

See Figure 1C for a visualization of these measures in an example neuron.

## 2.4 Selectivity Measurements

A neuron is selective to a task variable if its firing rate is significantly and reliably affected by the identity of that task variable. In this task, each condition contains three task variables: task type (recall or recognition), the identity of the first cue, and the identity of the second cue. Therefore, we used a 3-way ANOVA to determine if a given neuron’s firing rate was significantly (p<.05) affected by a task variable or combination of task variables. Selectivity can be of two types: pure or nonlinearly mixed (referred to as just “mixed”), based on which terms in the ANOVA are significant. If a neuron has a significant effect from one of the task variables, for example, it would have pure selectivity to that variable. Interaction terms in the ANOVA represent nonlinear effects from combinations of variables. Therefore, any neurons that have significant contributions from interaction terms as determined by the ANOVA have nonlinear mixed selectivity. As an example, if a neuron’s firing rate can be described by a function that is linear in the identity of the task type, the identity of the second cue, and the identity of the combination of task type and first cue, then that neuron has pure selectivity to task type (TT), pure selectivity to cue 2 (C2) and mixed selectivity to the combination of task type and cue 1 (TTxC1). Note that having pure selectivity to two or more task variables is not the same as having nonlinear mixed selectivity to a combination of those task variables.

We also investigate whether the nonlinear interactions we observe indicate supra- or sublinear effects. To do this we fit a general linear model that includes 2nd-order interaction terms to each neuron’s response. The signs of the coefficients for the 2nd-order terms indicate whether a certain nonlinear effect leads to a response higher (supralinear) or lower (sublinear) than expected from a purely additive relationship.

## 2.5 Clustering Measurement

Beyond the numbers of neurons selective to different task variables, an understanding of how preferences to task variable identities cluster can inform network models. For this, we use a method that is inspired by the Projection Angle Index of Response Similarity (PAIRS) measurement as described in Raposo et al. (2014). For this measure each neuron is treated as a vector in selectivity space, where the dimensions represent preference to a given task variable identity (Figure 1D). To get these values, neuronal responses are fit with a general linear model (GLM) to find which task variable identities significantly contribute to the firing rate. Note that this gives a beta coefficient for each value of each task variable, such as cue 1=B. These values dictate how the firing rate changes as task variable identities differ from the reference condition Task Type = Recognition, Cue 1 =A, and Cue 2 = B. Formally: *FR* = *FR*_*ref*_ + *β*_1_[*TT* = *Recall*] + *β*_2_[*C*1 = *B*]+ *β*_3_[*C*1 = *C*]+ *β*_4_[*C*1 = *D*]+ *β*_5_[*C*2 = *A*]+ *β*_6_[*C*2 = *C*]+ *β*_7_[*C*2 = *D*]. The beta values found for each cell via this method are shown in Figure 3C (non-significant coefficients — those with p>.05 — are set to 0).

This analysis does not include interaction terms (second- or third-order terms). The reason for this is partly that, given the relatively low number of trials, the high dimensional full GLM model would be difficult to confidently fit. In addition, analysis of clustering in a high-dimensional space (the full model would yield a 45-dimensional space) with a relatively small number of neurons would be difficult to interpret. Therefore, we look only at how the cells cluster according to their preference of the identities associated with the pure terms.

The coefficients derived from the GLM define a vector in a 7-D vector space for each neuron (see Figure 1D for a schematic). The clustering method compares the distribution of vectors generated by the data (each normalized to be unit length) to a uniform distribution on the unit hypersphere in order to determine if certain combinations of preferences are more common than expected by chance.

In PAIRS (Raposo et al., 2014), this comparison is done by first computing the average angle between a given vector and its k nearest neighbors and seeing if the distribution of those values differs between the data and a random population. That approach is less reliable in higher dimensions, therefore we use the Bingham test instead of PAIRS (Mardia and Jupp, 2000). The Bingham test calculates the test statistic 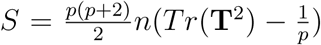. This statistic, which we refer to as the clustering value, measures the extent to which the scatter matrix, **T**, (an approximation of the covariance matrix) differs from the identity matrix (scaled by 1/*p*), where *p* and *n* are the dimensions of the selectivity space (7) and the number of cells (90), respectively. The higher this value is, the more the data deviates from a random population of vectors wherein selectivity values are IID. Thus, a high value suggests that neurons in the population cluster according to task variable identity preferences. In order to put this clustering value into context we compared the value found from the data to two distributions: one generated by shuffled data and one generated from data designed to be highly clustered. For the shuffled data, we created “fake” cell vectors by shuffling the selectivity values across all cells. For the clustered data, we created 3 categories of fake cells, each defined by pure selectivity to two specific task variable identities. A population of 90 cells was created by combining 30 cells from each category (the population was also designed to have the same average firing rate and *FF*_*T*_ of the data). This results in a population that has 3 clear clusters of cell types in selectivity space. 100 populations based on each type of fake data were created in order to generate distributions that represent random and clustered data.

Using the Gine-Ajne test of uniformity on the hypersphere (Giné, 1975) gives very similar results to the Bingham test results.

## 2.6 Circuit Model

To explore the circuit mechanisms behind PFC selectivity, we built a simple two-layer neural model, modeled off of previous work (Barak et al., 2013) (see Figure 5A for a diagram). The first layer consists of populations of binary neurons, with each population representing a task variable identity. To replicate a given condition, the populations associated with the task variable identities of that condition are turned on (set to 1) and all other populations are off (set to 0). Each population has a baseline of 50 neurons. To capture the biases in selectivities found in this dataset (particularly the fact that, in the 900ms period we used for this analysis, many more cells show selectivity to task type than cue 2 and to cue 2 than cue 1), the number of neurons in the task type and cue 2 populations are scaled by factors that reflect these biases (80 cells in each task type population and 60 in each cue 2 population). The exact values of these weightings do not have a significant impact on properties of interest in the model.

The second layer represents PFC cells. These cells get weighted input from a subset of the first layer cells. Cells from the input layer to the PFC layer are connected with probability .25 (unless otherwise stated), and weights for the existing connections are drawn from a Gaussian distribution (*μ*_*W*_ = .207, and *σ*_*W*_ = *μ*_*W*_ unless otherwise stated. Because negative weights are set to 0, the actual connection probability and *σ*_*W*_ may be slightly lower than given).

The activity of a PFC cell on each trial, *t*, is a sigmoidal function of the sum of its inputs:

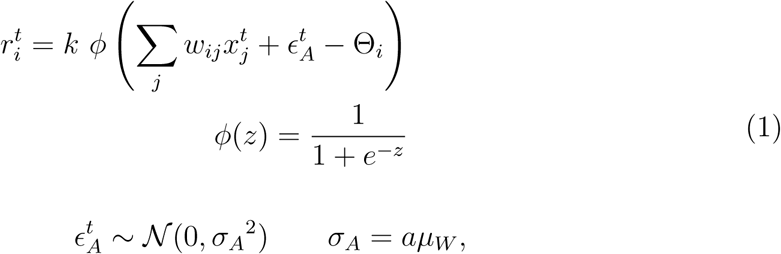

where *x*_*j*_ is the activity (0 or 1) of the *j*^*th*^ input neuron and *w*_*ij*_ is the weight from the *j*^*th*^ input neuron to the *i*^*th*^ output neuron. Θ_*i*_ is the threshold for the *i*^*th*^ output neuron, which is calculated as a percentage of the total weight it receives: Θ_*i*_ = *λ*Σ_*j*_*w*_*ij*_. The *λ* value is constant across all cells, making Θ cell-dependent. *k* scales the responses so that the average model firing rate matches that of the data.

Two sources of noise are used to model trial-to-trial variability. *∈*_*A*_ is an additive synaptic noise term drawn independently on each trial for each cell from a Gaussian distribution with mean zero. The standard deviation for this distribution is controlled by the parameter *a*, which defines *σ*_*A*_ in units of the mean of the weight distribution, *μ*_*W*_. The second noise source is multiplicative and depends on the activity of a given cell on each trial:

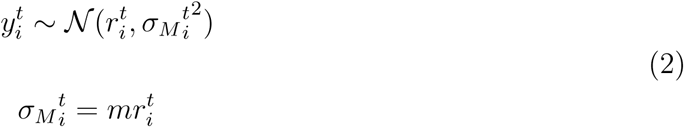

Thus, the final activity of an output PFC cell on each trial, 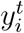, is drawn from a Gaussian with a standard deviation that is a function of 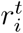. This standard deviation is controlled by the parameter *m*. Both *m* and *a* are fit to make the model *FF*_*T*_ match that of the data.

To make the model as comparable to the data as possible, ten trials are run for each condition and 90 model PFC cells are used for inclusion in the analysis.

## 2.7 Hebbian Learning

A simplified version of Hebbian learning is implemented in the network in a manner that captures the “rich get richer” nature of Hebbian learning while keeping the overall input to an individual cell constant. In traditional Hebbian learning, weight updates are a function of the activity levels of the pre- and post-synaptic neurons: Δ*w*_*ij*_ = *g*(*x*_*j*_*, y*_*i*_). In this simplified model we use connection strength as a proxy for joint activity levels: Δ*w*_*ij*_ = *g*(*w*_*ij*_). We also implement a weight normalization procedure so that the total input weight to a cell remains constant as weights change.

To do this, we first calculate the total amount of input each output cell, *i*, receives from each input population, *p*:

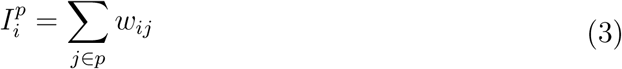

The input populations (each corresponding to one task variable identity) are then ranked according to this value. The top *N*_*L*_ populations according to this ranking (that is, those with the strongest total weights onto to the output cell) have the weights from their constituent cells increased according to:

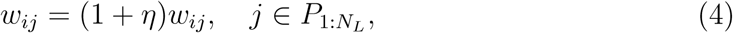

where *η* is the learning rate (set to .2 unless otherwise stated). This amounts to a multiplicative scaling of synaptic weights, which is compatible with experimental observations (Loewenstein et al., 2011; Turrigiano et al., 1998). After this, all weights into the cell are normalized via:

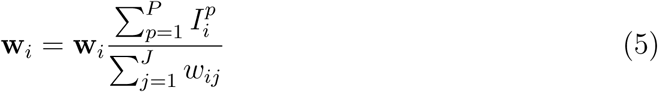

Note, the numerator in the second term is the sum of all weights into the cell before Eqn. 4 is applied and the denominator is the sum after it is applied. As learning progresses according to this rule, weights from cells that aren’t in the top *N*_*L*_ populations trend to zero. At that point, each learning step increases the weights of all remaining connections by *η* and normalizes them all by the same amount, resulting in no further changes in the weight matrix.

In this work, two versions of Hebbian learning are tested. In the unrestricted, or “free”, learning condition described above, the top *N*_*L*_ populations are chosen freely from all input populations (equivalently, all task variable identities) based solely on the total input coming from each population after the random weights are assigned. The alternative, “constrained” learning, is largely the same, but with a constraint on how these top *N*_*L*_ populations are chosen: all task variables must be represented before any can be repeated. So, two populations representing different identities of the same task variable (e.g., cue 1 A and cue 1 B) will not both be included in the *N*_*L*_ populations unless both other task variables already have a population included (which would require that *N*_*L*_ > 3). So, with *N*_*L*_ = 3, exactly one population from each task variable (task type, cue 1, cue 2) will have weights increased. This variant of the learning procedure was designed to ensure that inputs could be mixed from different task variables, to increase the likelihood that mixed selectivity would arise. Both forms of learning are demonstrated for an example cell in Figure 5B.

In both forms of learning, the combination of weight updating and normalization is applied to each cell once per learning step.

## 2.8 Classification Performance

The measures of selectivity we have looked at in the data are important for the ability of a population to represent task information in a way that can be readily readout. We also test directly the ability to readout task information from our model populations using linear discriminant analysis (LDA). We generate 20 trials per condition from the model and use 10 to train the classifiers and 10 to test. Three separate classifiers are trained to read out each of the three linear terms: task type identity, cue 1 identity, and cue 2 identity. The average performance across these three tasks gives the “linear” performance. An additional four classifiers were trained to read out each of the joint identities of task type-cue 1, task-type 2, cue 1-cue 2, and task type-cue 1-cue 2. The average performance across these four tasks is called the “higher order” performance.

We also conduct an explicit test of the model’s ability to perform a non-linearly separable task. For this, all combinations of identities for cue 1 and cue 2 are generated as inputs to the network, and the classification task is to determine if the identities are the same or different (the task type input is held constant). Fifty trials are used for training (using LDA) and fifty for testing. We also measure the ability of the input population to perform this task (by using the binary input population activity directly), in which case additive noise is used to generate multiple trials, and the mean firing rate and *FF*_*T*_ are fit to match that of the data.

## 2.9 Toy Model Calculations

To make calculations and visualizations of the impacts of learning easier, we use a further simplified toy model (see Figure 8A (left) for a schematic). Instead of a sigmoidal nonlinearity, the heaviside function is used. The toy model has two task variables (T1 and T2) and each task variable has two possible identities (A or B). Four random weights connect these input populations to the output cell: *W*_1*A*_*, W*_1*B*_*, W*_2*A*_*, W*_2*B*_. On each condition, exactly one task variable identity from each task variable is active (set to 1). This gives four possible conditions, each of which is plotted as a point in the input space in Figure 2. The threshold is denoted by the dotted lines. If the weighted sum of the inputs on a given condition is above the threshold, the cell is active (green), otherwise it is not.

**Figure 2:**
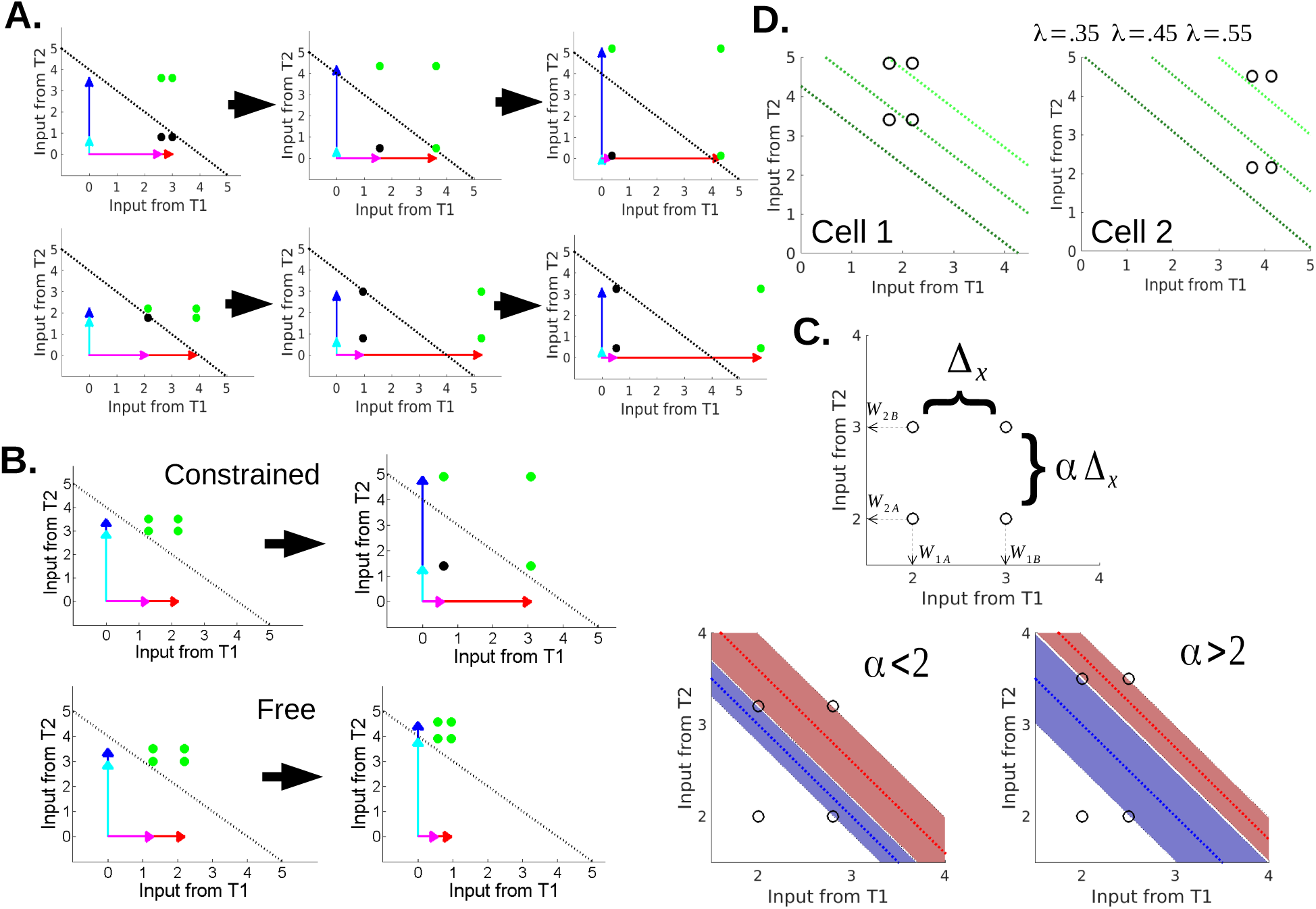
Signal and noise representation for the toy model shown in Figure 8A. Strength of weights from the 4 input populations are given as arrows in (A and B) and the threshold for the heaviside function is shown as a dotted line. The cell is active for conditions above the threshold (green). Weight arrows omitted for visibility in (C and D). A.) Learning causes the representation of conditions to change. This can change selectivity in multiple ways. Shown here: pure selectivity turns into mixed selectivity (top) and mixed selectivity turns into pure (bottom). B.) Constrained and free learning can lead to different signal changes. Constrained learning (top) guarantees that one population from each task variable is increased. This ensures that the representation spreads out. In this case, the cell goes from no selectivity to mixed selectivity. With these starting weights, free learning increases both populations from T2, and the cell does not gain selectivity. C.) Noise robustness can be thought of as the range of thresholds that can sustain a particular type of selectivity. Relative noise robustness of mixed and pure selectivity depends on the shape of the representation. *α* is the ratio of the differences between the weights from each task variable (top). In the two figures on the bottom, blue (red) dotted lines show optimal threshold for pure (mixed) selectivity and shaded areas show the range of thresholds created by trialwise additive noise that can exist without altering the selectivity. When *α <* 2, mixed selectivity is robust to larger noise ranges (bottom left). When *α >* 2, pure selectivity is more robust (bottom right). Given normally-distributed weights, *α >* 2 is more common. D. Two example cells showing how selectivity changes with changing *λ*. Sets of weights for both cells are drawn from the same distribution. The resulting thresholds at 3 different *λ* values (labeled on the right cell but identical for each) are shown for each cell. With the smallest *λ*, neither example cell has selectivity. With the middle *λ* value Cell 1 gains mixed. Cell 2 gains pure selectivity, which it retains at the higher *λ*, while Cell 1 switches to the other type of mixed

**Figure 3:**
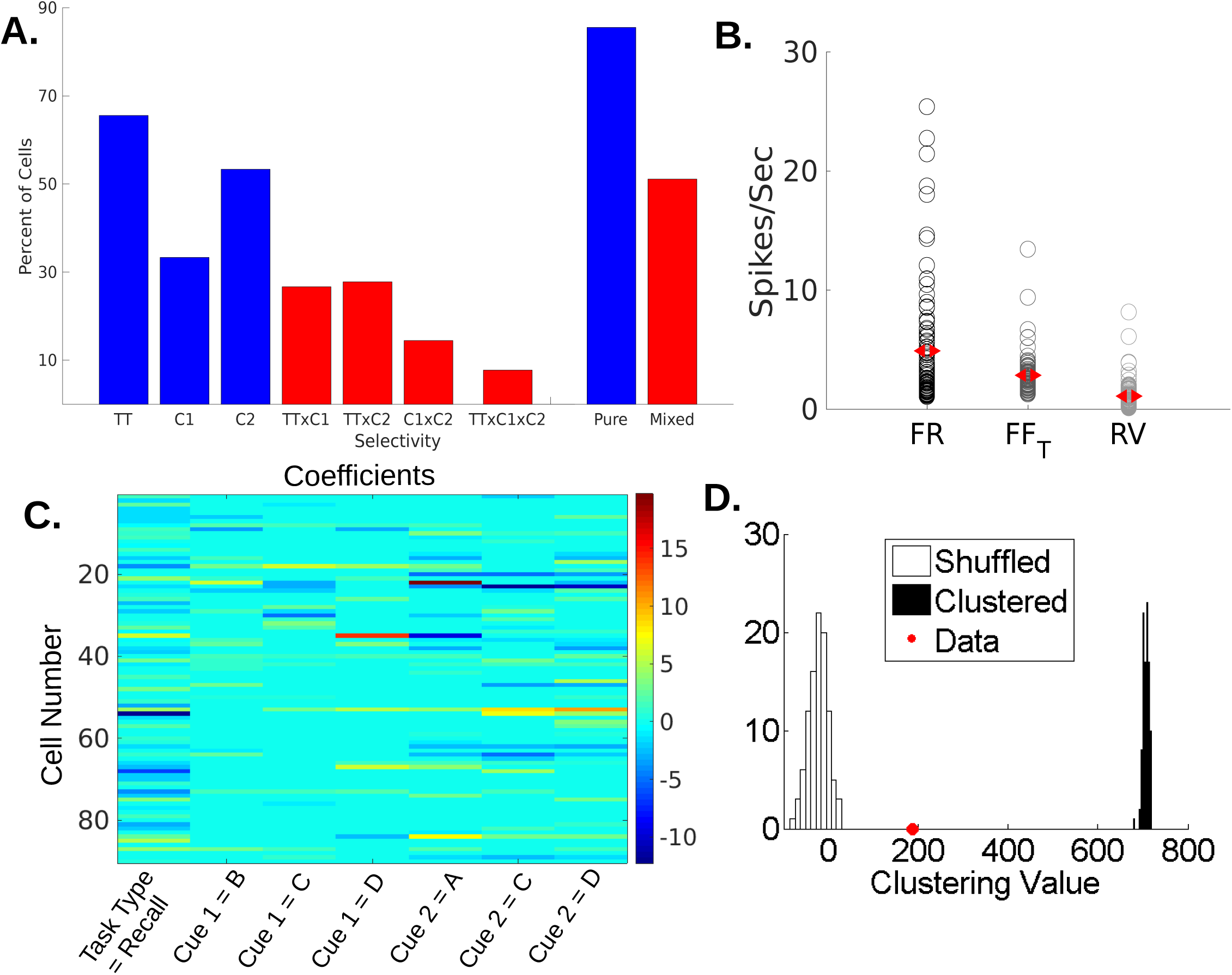
Results from the experimental data. A.) Selectivity profile of the 90 cells analyzed. A cell had pure selectivity to a given task variable if the term in the ANOVA associated with that task variable (TT=Task Type, C1=Cue 1, C2=Cue 2) was significant (p<.05). A cell had nonlinear mixed selectivity to a combination of task variables if the interaction term for that combination (TTxC1=Task Type × Cue 1, TTxC2=Task Type × Cue 2, C1xC2=Cue 1 × Cue 2, TTxC1xC2=Task Type × Cue 1 × Cue 2) was significant. On the right of the vertical bar are the percent of cells that had at least one type of pure selectivity (blue) and percent of cells that had at least one type of mixed selectivity (red). B.) Values of firing rate, *FF*_*T*_, and RV for this data. Each open circle is a neuron and the red markers are the population means. C.) Beta coefficients from GLM fits for each cell. The condition wherein Task Type = Recognition, Cue 1 = A, and Cue 2 = B was used as the reference condition. These values were used to determine the clustering value D.) Clustering values for data and comparison populations. The red dot shows the clustering value calculated using the GLM coefficients from the data. The shuffled data comes from shuffling the GLM coefficients across cells. The clustered data derives from populations of fake cells designed to have 3 different categories of cell types defined according to selectivity.

**Figure 4:**
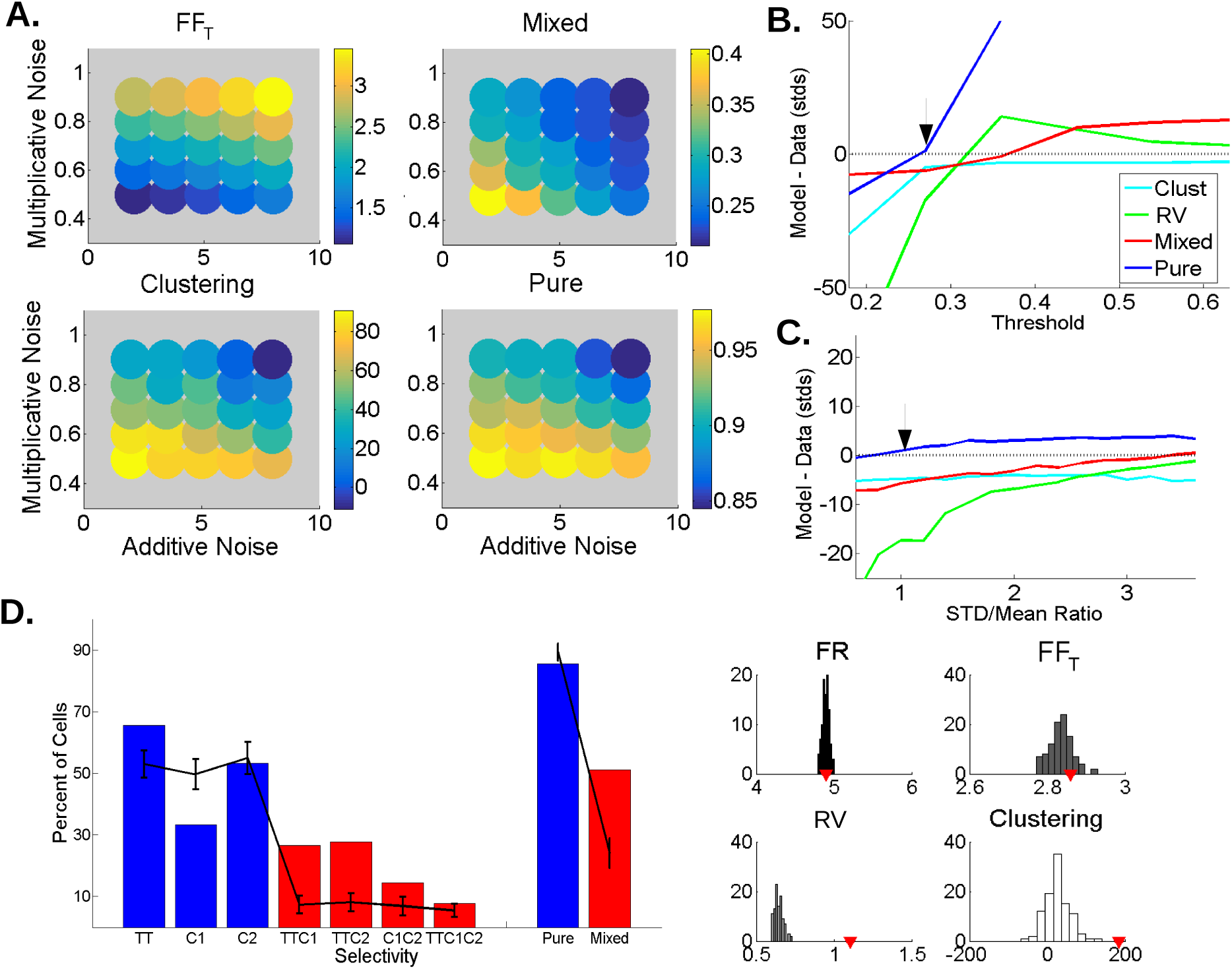
Results from the model without learning. A.) *FF*_*T*_ and other measures can be controlled by the additive and multiplicative noise parameters. Each circle’s color shows the value for the given measure averaged over 25 networks, for a set of *a* and *m* values (see Methods). *FF*_*T*_ scales predictably with both noise parameters. Fraction of cells with mixed selectivity, fraction of cells with pure selectivity, and clustering scale inversely with the noise parameters. Other model parameters are taken from the arrow locations in (B) and (C). B.) How the threshold parameter, *λ*, affects measures of selectivity. Lines show how the average value of the given measure in the model (in units of standard deviations calculated over 100 random instantiations of the model) differs from the data as a function of the threshold parameter *λ*, where Θ_*i*_ = *λ*Σ_*j*_*w*_*ij*_ At each point noise parameters are fit to keep *FF*_*T*_ close to the data value. Note that std values for mixed selectivity and clustering remain steady across threshold values at approximately 4% and 20.7 respectively. RV std however increases from .0087 to 4.3 spikes/sec and pure selectivity std trends toward zero as all cells gain pure selectivity. C.) Same as (B), but varying the width of the weight distribution rather than the threshold parameter. Here, RV std increases only slightly, from .02 to .048 spikes/sec, pure selectivity std decreases slightly from 4.0% to 2.5% and mixed selectivity and clustering stds remain fairly constant around 4.9% and 31.2 respectively. D.) Example of the model results at the points given by the black arrows in (B) and (C). On the left, blue and red bars are the data values as in Fig 2. The lines are model values (averaged over 100 networks, errorbars *±*1 std). On the right, histograms of model values over 100 networks. The red markers are data values. This model has no learning.

**Figure 5:**
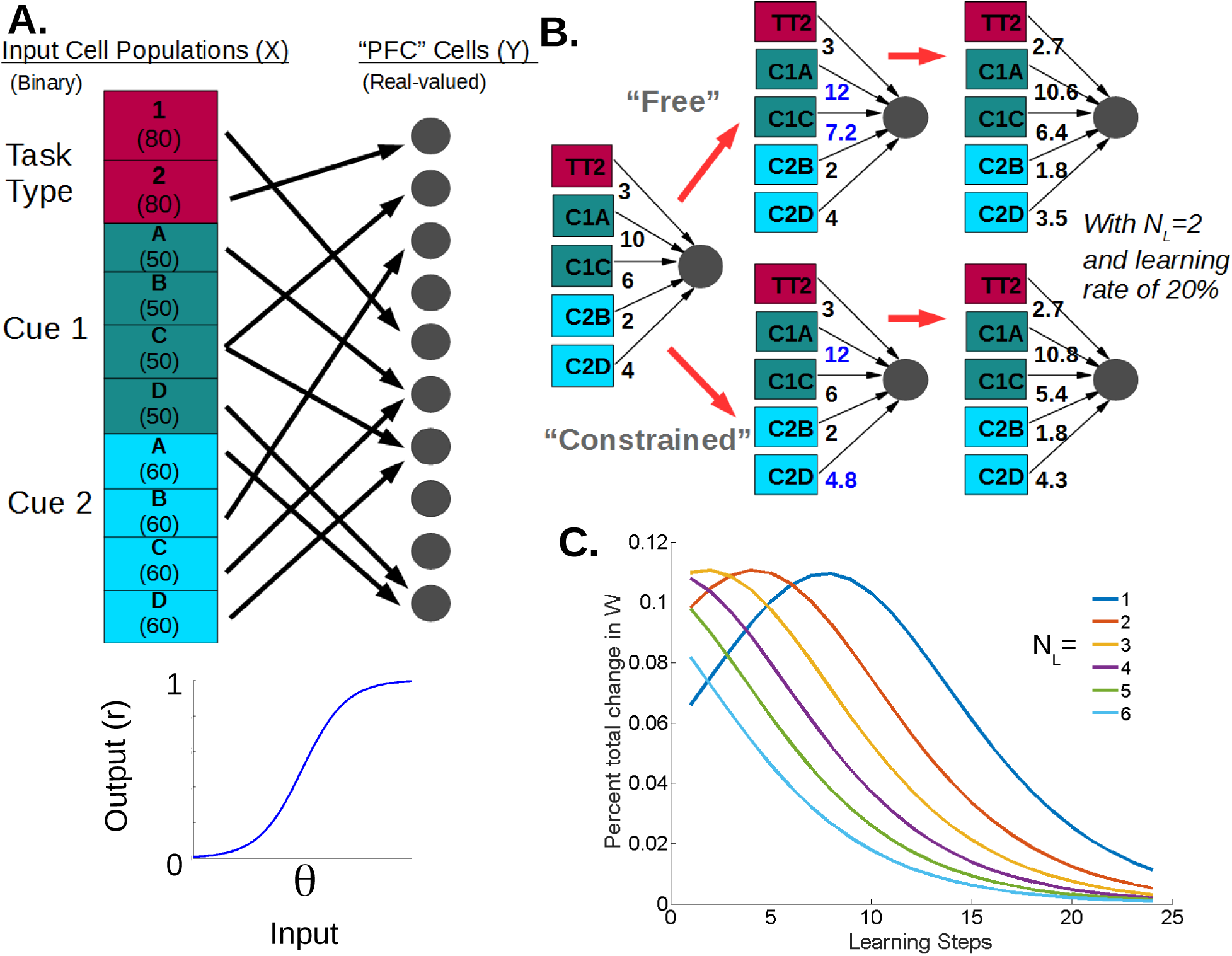
The full model and how learning occurs in it. A.) The model consists of groups of binary input neurons (colored blocks) that each represent a task variable identity. The number of neurons per group is given in parenthesis. Each PFC cell (gray circles) receives random input from the binary cells. Connection probability is 25% and weights are Gaussian-distributed and non-negative. The sum of inputs from the binary population and an additive noise term are combined as input to a sigmoidal function (bottom). The output of the PFC cell on a given trial is a function of the output of the sigmoidal function, *r* and a multiplicative noise term (see Methods). The threshold, Θ, is given as percentage of sum total of the weights into to each cell B.) Two styles of learning in the network, both of which are based on the idea that the input groups that initially give strong input to a PFC cell have their weights increased with learning (sum of weights from each population are given next to each block). In free learning, the top *N*_*L*_ input populations are chosen freely. In this example, that means two groups from the cue 1 task variable have their weights increased (marked in blue). In constrained learning, the top *N*_*L*_ populations are chosen with the constraint that they cannot come from the same task variable. In this case, that means that cue 2D is chosen over cue 1C despite the latter having a larger summed weight. In both cases, all weights are then normalized. C.) Learning curves as a function of learning steps for different values of *N*_*L*_. Strength of changes in the weight matrix expressed as a percent of the sum total of the weight matrix are plotted for each learning step (a learning step consists of both the weight increase and normalization steps). Different colors represent different *N*_*L*_s.

**Figure 6:**
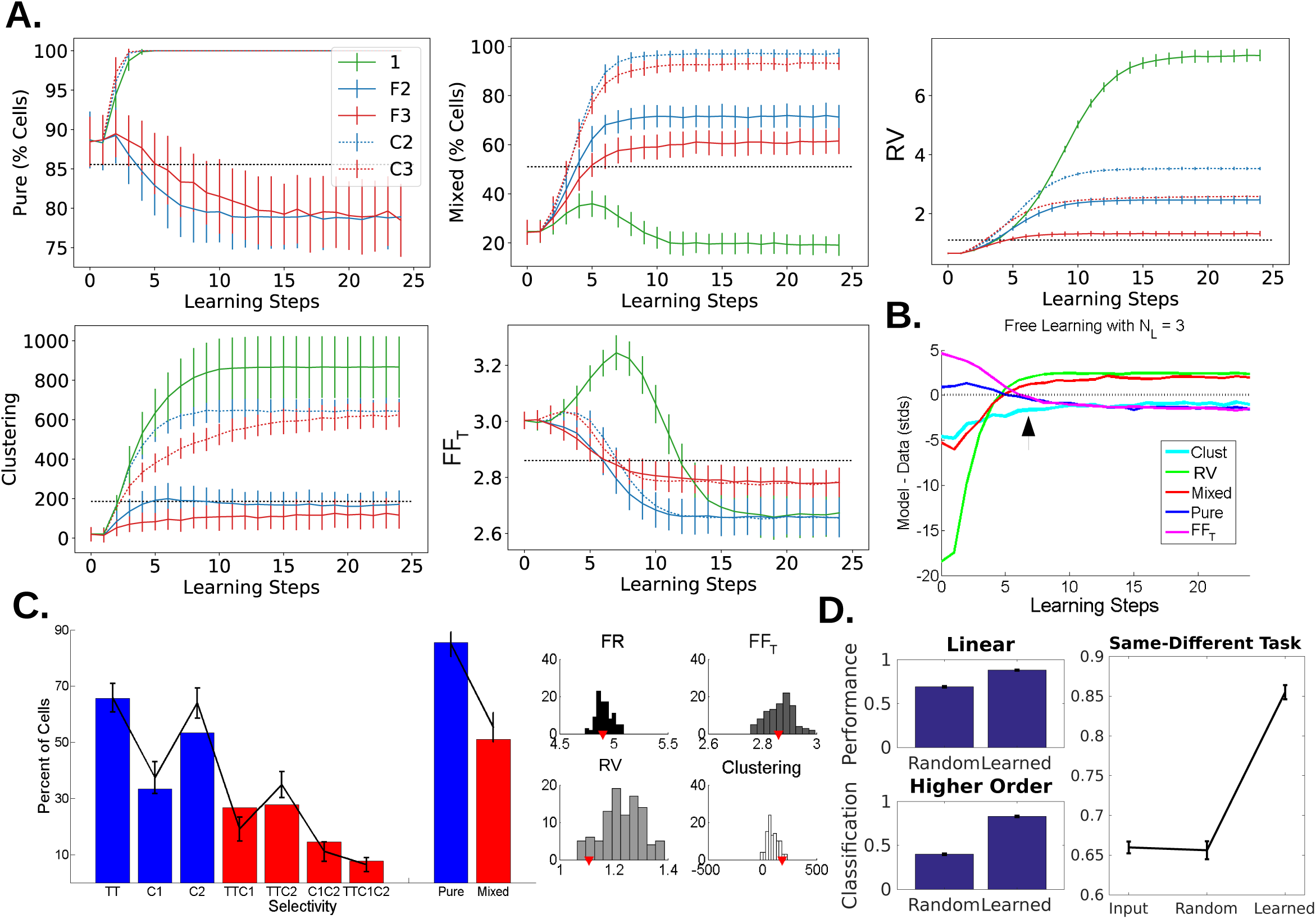
The model with learning. A.) How selectivity measures change with learning. In each plot, color represents *N*_*L*_ value, solid lines are free learning, and dotted lines are constrained learning (only one line is shown for *N*_*L*_ = 1 as the free and constrained learning collapse to the same model in this circumstance). Step 0 is the random network. Black dotted lines are data values and errorbars are *±*1 std over 100 networks. In the pure selectivity plot, with constrained learning and when *N*_*L*_ = 1, the value maxes out at 100% in essentially all networks, leading to vanishing errorbars. B.) All measures as a function of learning for the *N*_*L*_ = 3 free learning case. Values are given in units of model standard deviation away from the data value as in Figure 4B and C. C.) The model results at the learning step indicated with the black arrow in (B). On the left, blue and red bars are the data values as in Figure 3. The lines are model values (averaged over 100 networks, errorbars *±*1 std). On the right, histograms of model values over 100 networks. The red markers are data values. Here, the model provides a much better match to the data. D.) Decoding performance increases with learning. Average performance of classifiers trained to readout linear terms (top left) and higher order terms (bottom left) from PFC population activity increases after learning compared to the random network (learned model indicted by arrow in (B)). Errorbars are *±*1 SEM, over 10 random instantiations of the network. Read out of same vs. different cue identities is better when using the PFC population after learning (right).

**Figure 7:**
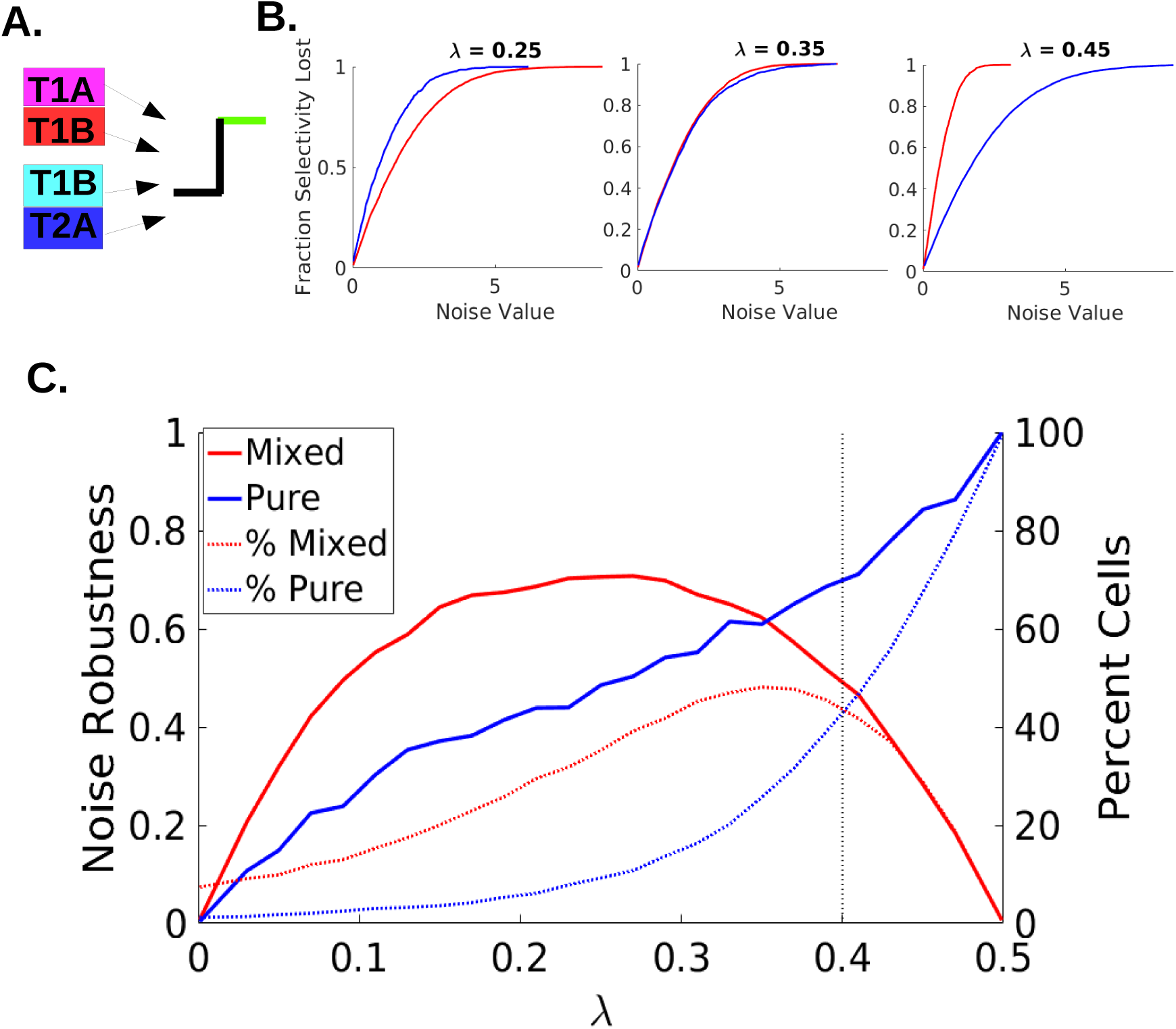
How noise robustness varies with threshold in a random network using the toy model A.) Schematic of the toy model: four input populations (two from each task variable) send weighted inputs to a cell with a threshold (Θ) nonlinearity B.) For a given noise value, the fraction of cells that would lose selectivity if that noise value were used. Values are separated for cells with pure (blue) and mixed (red) selectivity. Three λ values shown, where Θ = λΣW. C.) Based on plots like those in (B), the noise value at which 50% of cells have lost selectivity is calculated (”Noise Robustness” refers to these values normalized by the peak value. Higher values are better) and plotted as a function of λ (solid lines). On the same plot, the percent of cells with each type of selectivity in the absence of noise is shown (dotted lines). The black doted line marks a λ value at which the probability of mixed and pure selective cells is equal, but their noise robustness is unequal. This plot is mirror-symmetric around λ = .5

**Figure 8:**
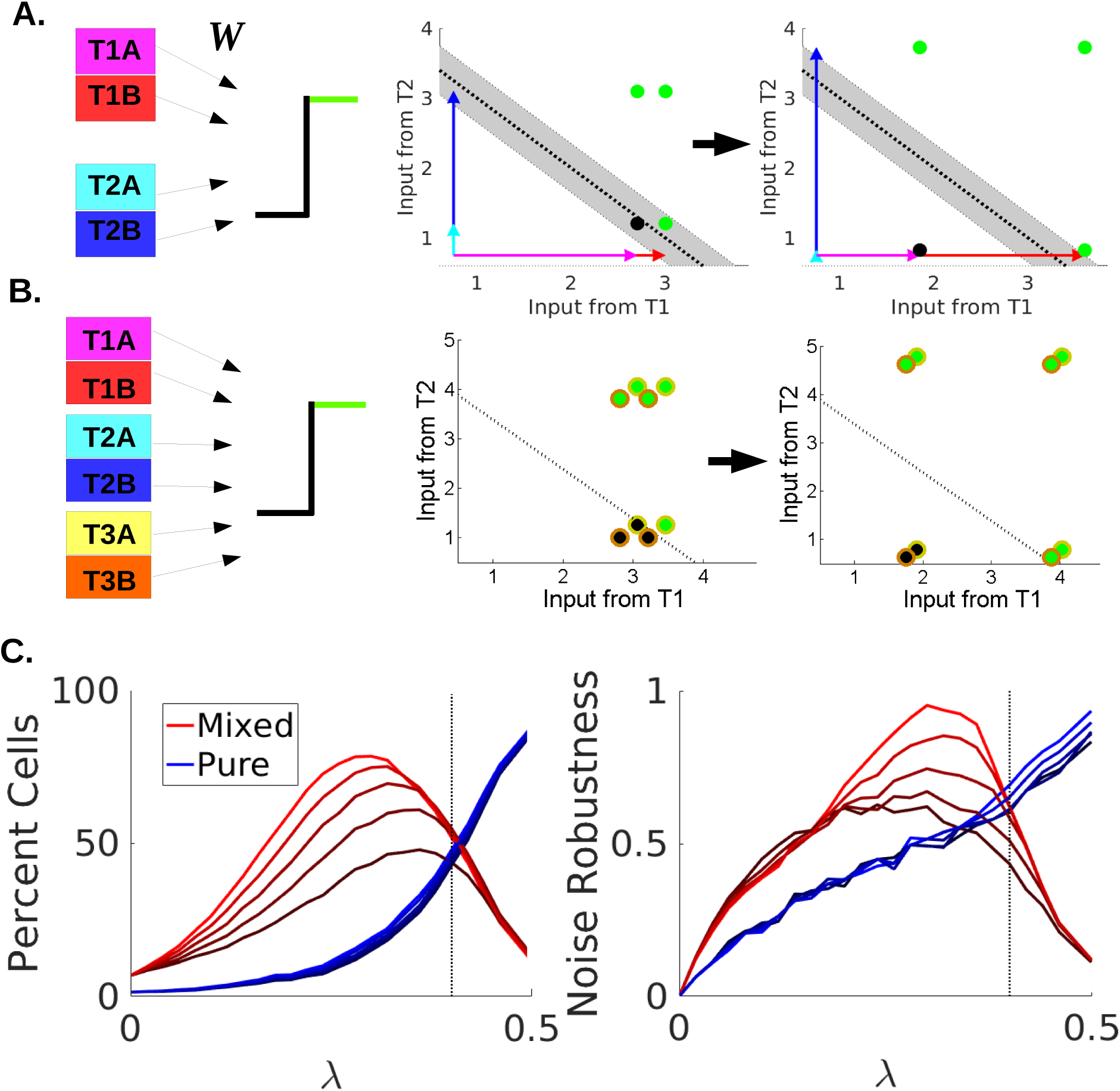
How learning impacts noise robustness A.) A simple toy cell (left) with 2 task variables is used to w the effects of learning. The 4 possible conditions are plotted as dots (green if above threshold, black if, with the threshold as a dotted black line. Colored arrows represent the weights from each population. ore learning (middle), the cell’s input on two of the conditions falls within the range of the shifting threshold ted by additive noise (gray area). After learning, all conditions are outside the noise range. B.) A third variable is added to the model and is another source of additive noise from the perspective of T1-T2 ctivity. The model’s outputs are color-coded according to which T3 population is active. Weight arrows omitted for visibility. After learning with *N*_*L*_ = 2, input strength from T3 populations are decreased and points from the same T1-T2 condition are closer together (less noisy). C.) How the percent of cells with a n selectivity (left) and their noise robustness (right) change with constrained learning as a function of the shold parameter *λ*. Learning steps are symbolized by increasing color brightness (the darkest line is the dom model as displayed in Figure 7C, and the dashed line shows where the percent of mixed and pure are same in the random model)

The toy model follows the same learning rules defined for the full model. Examples of the impacts of learning on the representation of the 4 conditions are seen in Figure 2A and B.

A cell’s selectivity is more robust to additive noise (which functions like a shift in threshold) if there is a large range of threshold values for which its selectivity doesn’t change. To explore noise robustness in this model, we will define:

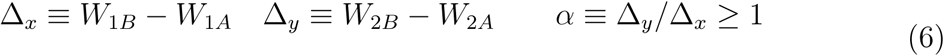

Thus, *α* is the ratio of the side lengths of the rectangle formed by the four conditions (see Figure 2C, top). Without loss of generality, we define the larger of the two sides as associated with T2, *W*_2*B*_ > *W*_2*A*_, and *W*_1*B*_ > *W*_1*A*_.

For the cell to display pure selectivity to T2, the following inequality must hold:

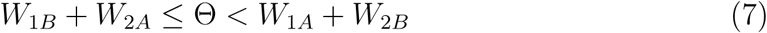

Therefore the range of thresholds that give rise to pure selectivity is:

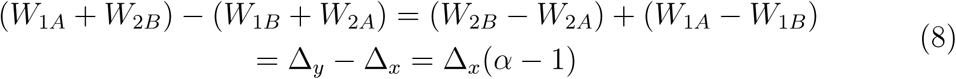

The analogous calculations for mixed selectivity (assuming the T1B-T2B condition is active only, but results are identical for T1A-T2A being the only inactive condition) are:

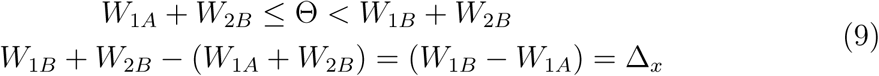

Thus, pure selectivity is more noise robust than mixed selectivity when *α >* 2. This imbalance can be seen in Figure 2C.

Now we show that, given weights drawn at random from a Gaussian distribution, *α >* 2 is more common than *α <* 2. The argument goes as follows: because Δ_*x*_ and Δ_*y*_ are differences of normally distributed variables, they are themselves normally distributed (with *μ* = 0, *σ* = 2*σ*_*w*_). The ratio of these differences is thus given by a Cauchy distribution. However, because *α* represents a ratio of lengths, we are only interested in the magnitude of this ratio, which follows a standard half-Cauchy distribution. Furthermore, *α* is defined such that the larger difference should always be in the numerator. Thus,

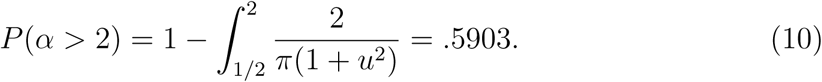

Therefore, the majority of cells can be expected to have *α >* 2 with random weights and thus higher noise robustness for pure selectivity than for mixed.

This comparison of noise robustness, however, assumes the threshold is placed at the most noise robust location for each type of selectivity. Here, the threshold is defined as a fraction of the total weight going into the cell: Θ = *λ*Σ*W*. As we increase *λ* then, the threshold is a line with slope of -1 that moves from the bottom left corner up to the top right. Examples of how this impacts selectivity are shown in Figure 2D. To investigate how noise robustness changes with *λ*, we generate a large (10000) population of cells, each with four random input weights drawn from a Gaussian with positive mean and constrained to be non-negative (qualitative results hold for many weight/variance pairs), and calculate the size of the additive noise shift needed to cause each cell to lose its selectivity (whichever it has).

Assuming a fixed threshold, we then explore how noise robustness varies with learning. In the case of constrained learning with *N*_*L*_ = 2, Δ_*x*_ and Δ_*y*_ both increase. According to Eqn. 7 and Eqn. 9, robustness to both selectivities increases with Δ_*x*_. The relative increase in robustness will depend on how *α* changes. It can be shown that if 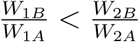 then Δ_*x*_ will expand more than Δ_*y*_ and *α* will decrease, meaning the increase in noise robustness favors mixed selectivity. If 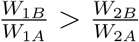, then *α* will grow, and the increase in noise robustness will be larger for pure than mixed. However, this condition is less common.

When *N*_*L*_ = 1, learning ultimately leads to a larger ratio between the side lengths. This is straightforward for *W*_2*B*_ > *W*_1*B*_ (Δ_*y*_ grows and Δ_*x*_ shrinks). However, if *W*_1*B*_ > *W*_2*B*_, *α* will first decrease as Δ_*x*_ grows and Δ_*y*_ shrinks. This is good for mixed noise robustness. The ratio then flips (Δ_*x*_ > Δ_*y*_), and Δ_*y*_ (the side that is now shorter) is still shrinking and Δ_*x*_ is growing. In this circumstance, if Δ_*y*_/Δ_*x*_ becomes less than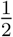, the representation will favor pure noise robustness over mixed. This flipping of *α* is possible for some cells when 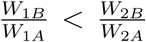 but the weights would likely plateau before *α* became less than 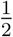, and so the drop in mixed selectivity does not occur.

In free learning with *N*_*L*_ = 2, cells that have *W*_1*A*_ > *W*_2*B*_, will see both weights from T1 increase and (due to the weight normalization) both weights from T2 decrease. Because the weights change in proportion to their value, Δ_*x*_ increases, Δ_*y*_ decreases and so *α* goes down. This leads to more noise robustness for mixed and less for pure. If *W*_2*A*_ > *W*_1*B*_, these trends are reversed and the cell has more noise robustness for pure and less for mixed.

## 2.10 Experimental Design and Statistical Analysis

As described in the Selectivity Measurements subsection above, the main statistical test used in this work was a 3-way ANOVA (within-subjects, with a total 23 degrees of freedom). Each of the 90 cells used had 10 trials from each condition. As part of calculating the clustering value (see Clustering Measurement subsections above), we calculated the p-value for the F statistic of the hypothesis test that each coefficient in our General Linear Model was equal to 0. All analyses were performed in MATLAB.

## 3. Results

In this study, we analyzed various measures of selectivity of a population of PFC cells recorded as an animal carried out a complex delayed match-to-sample task. Through this process, several properties of the representation in PFC were discovered and a simple circuit model that included Hebbian learning was able to replicate them. These properties, combined with the modeling results, provide support for the notion that PFC selectivities are the result of Hebbian learning in a random network.

## 3.1 PFC Population is Moderately Specialized and Selective

The average firing rate of cells in this population was 4.9 *±* 5.1 spikes/sec. Fano Factor analyses provided measurements of the noise and density of response in the data (Figure 3B). The average value of the across-trial Fano Factor (*FF*_*T*_ = 2.8 *±* 1.7 spikes/sec), shows that the data has elevated levels of noise compared to a Poisson assumption. Looking at response variability (RV)—a measure of how a cell’s response is distributed across conditions—suggests that PFC cells are responding densely across the 24 conditions (*RV* = 1.1 *±* 1.1 spikes/sec, for comparison, at the observed average firing rates, a cell that responded only to a single condition would have *RV*≈ 120, one that responded to two conditions would have *RV*≈ 57). This finding suggests that these cells are not responding sparsely and are not very specialized for the individual conditions of this task.

Each condition is defined by a unique combination of 3 task variables: task type, identity of image cue 1 and identity of image cue 2 (Figure 1A). Selectivity to task variables was determined via a 3-way ANOVA. The results of this analysis are shown in Figure 3A. This figure shows the percentage of cells with selectivity to each task variable and combination of task variables (as determined by a significant (p<.05) term in the ANOVA). A cell that has selectivity to any of the regular task variables (task type, cue 1, cue 2) has pure selectivity, while a cell that has selectivity to any of the interaction terms (combination of task variables such as task type × cue 1, task type × cue 2, etc) has nonlinear mixed selectivity. The final two bars in Figure 3A show the number of cells with pure and mixed selectivity defined this way. Note that a cell can have both pure and mixed selectivity, thus the two values sum to more than 100%.

The majority of cells (77/90) showed pure selectivity to at least one task variable. But the population shows clear biases in the distribution of these pure selectivities: task type selectivity is the most common (59 cells) and cue 2 is represented more than cue 1 (48 vs. 30 cells) (these biases are observable in the GLM fits as well, see Figure 3C). This latter effect may be due to the time at which these rates were collected: these rates were taken during the second delay, which comes directly after the presentation of the second cue. The former effect is perhaps more surprising. While the task type is changed in blocks and thus knowable to the animal on each trial (with the exclusion of block changes), there is no explicit need for the animal to store this information: the presence of a second sequence or an array of images will signal the task type without the need for prior knowledge. However, regardless of its functional role in this task, contextual encoding is a common occurrence (Eichenbaum et al., 2007; Komorowski et al., 2013). Furthermore, the fact that the recall task is more challenging than the recognition task may contribute to clear representation of task type. That is, it is possible that the animals keep track of the task type in order to know how much effort to exert during the task.

Approximately half of the cells (46) had some form of mixed selectivity, mostly to combinations of two task variables. The population had a roughly equal balance of both supra- and sublinear effects of these 2-way interactions (ratio of positive to negative terms: 1.07). The small number of cells with selectivity to the 3-way interaction term (TTxC1xC2) is consistent with the relatively low value of RV in this population, as a strong preference for an individual condition would lead to a high RV. The number of cells with only mixed selectivity was low (only 1 out of 90 cells), 32 cells had only pure selectivity, and 12 cells had no selectivity.

We use a population-level analysis inspired by (Raposo et al., 2014) to measure the extent to which cell types are clustered into categories. Here, we used this analysis to determine if cells cluster according to their responsiveness to different task variable identities (i.e., recognition vs recall). That is, are there groups of neurons which all prefer the same task type and image identities, beyond what would be expected by chance? In order to explore this, we first use a general linear model (GLM), with task variable identities as regressors, to fit each neuron individually. The beta coefficients from these fits define a neuron’s position in selectivity space (these beta coefficient values, which represent how the identity of each task variable changes a neuron’s firing rate as compared to the reference condition, are shown in Figure 3C. A schematic of how the clustering measure works is shown in Figure 1D). After normalizing each vector, the clustering measure then determines the extent to which the population of vectors deviates from a uniform distribution on the unit hypersphere. The data had a clustering value of 186.2. Comparing this to the mean values of two distributions of artificially generated populations suggests the data has a mild but significant deviation from random: the average clustering value for populations generated by randomly shuffling the coefficient values is -23*±*22, and the average value of populations that have 3 distinct clusters of selectivity is 706.7*±*6.8. As the data clustering value sits in between these values and closer to the shuffled data, we conclude that some structure does exist in the data, yet the cells in this population do not appear to form strongly separable categories as defined by task variable identity preference (Figure 3D).

## 3.2 Circuit Model without Hebbian Learning Cannot Replicate Mix of Density and Specialization

A simple circuit model was made to replicate the selectivity properties found in the data. The model contains two layers: an input layer consisting of binary neurons that represent task variable identities and an output layer consisting of “PFC” neurons which get randomly-weighted input from the first layer and whose activity is a nonlinear function of the sum of that input. The model also has two forms of noise: an additive term applied before the nonlinearity (which replicates input/background noise, and implicitly shifts the threshold of the cell), and a multiplicative term applied after (which enforces the observed relationship between firing rate and variance) (see Methods and Figure 5A).

The output of the initial circuit model, prior to any Hebbian learning, was analyzed in the same way as the data to determine if it matched the properties found in PFC. The results of this can be found in Figure 4. First, in Figure 4A, we demonstrate the impact of the noise parameters on *FF*_*T*_, pure and mixed selectivity, and the clustering value. As expected, increasing the additive and/or multiplicative noise terms increases the *FF*_*T*_, as this is a measure of trial variability. Increasing within-condition noise also makes it less likely that a cell will show significant differences across conditions, and thus the percentage of cells with pure and mixed selectivity are inversely related to the noise parameters, (the relative sensitivities of mixed and pure selectivity to noise will be discussed in depth later). For similar reasons, the clustering value also decreases with noise (finding significant deviations from a uniform distribution is less likely if cells do not show sufficiently strong preferences).

To determine the impact other properties of the model had on our measures of interest, we varied several other parameters. Figure 4B shows what happens at different values of the threshold parameter. Here, the threshold is given as the amount of input the cell needs to reach half its maximal activity, expressed as a fraction of its total input weight (keep in mind that, given the number of input cells in each population and the task structure, roughly one-third of input cells are on per trial). The colored lines are, for each measure, the extent to which the model differs from the data, expressed in units of the model’s standard deviation (calculated over 100 instantiations of the model). Due to the impact of noise parameters discussed above, at each point in this graph the noise parameters were fit to ensure the model was within *±* 1.5 standard deviations of the data *FF*_*T*_ (this generally meant that it varied from *∼*2.8 to 2.9).

With an increasing threshold, the RV (green line in Figure 4B) increases. This is because higher thresholds mean cells respond to only a few combinations of input, rather than responding similarly to many, and the RV is a measure of variability in response across conditions (note that while RV appears to peak at ≈.35 and decrease, this particular trend is driven by an increase in RV standard deviation; the mean continues to increase). The percentage of cells with mixed selectivity (red line) also increases with threshold. With a higher threshold, the majority of conditions give input to the cell that lies in the lower portion of the sigmoidal function (bottom of Figure 5A). The nonlinearity is strong here—with some input producing little to no response—thus, more cells can attain nonlinear mixed selectivity. Pure selectivity also increases with threshold, and the percent of cells with pure selectivity goes quickly to 100 (and the standard deviation of the model gets increasingly small). We go into more detail about the reliance of selectivity on threshold later.

The clustering value relies on cells having preference for task variable identities and so increases as selectivity increases initially. However, just having selectivity is not enough to form clusters, and so the clustering value in the model levels off below the data value even as the number of cells with pure selectivity reaches full capacity. Thus, with the exception of the clustering value, the model can reach the values found in the data by using different thresholds. As Figure 4B shows, however, at no value of the threshold are all measures of PFC response in the model simultaneously aligned with those in the data.

Figure 4C shows how the same measures change when the width of the weight distribution from input to PFC cells is varied. Here, the standard deviation of the distribution from which connection strengths are drawn (*σ*_*W*_) is given as a factor of the mean weight, *μ*_*W*_. Increasing this value increases pure and mixed selectivity as well as RV. Because a wider weight distribution increases the chances of a very strong weight existing from an input cell to an output cell, it makes it easier for selectivity to emerge (that is, the output cell’s response will be strongly impacted by the task variable identity the input cell represents). The RV increase occurs for similar reasons: a cell may have uneven responses across conditions due to strong inputs from single input cells. Clustering values, however, are unaffected by this parameter. At no point, then, can the model recreate all aspects of the data by varying the weight distribution. Furthermore, while values of mixed selectivity and RV approach the data values with large *σ*_*W*_/*μ*_*W*_, such large values are likely unrealistic. Data show that a *σ*_*W*_/*μ*_*W*_ ratio of around 1 is consistent with observations of synaptic strengths from several brain areas (Barbour et al., 2007).

Varying other parameters such as the mean weight, number of cells per population, and connection probability similarly doesn’t allow the model to capture all properties of the data (not shown).

Figure 4D shows the values of the model as compared to the data for the set of parameters marked with arrows in Figure 4B and 4C. For reasons that will be discussed more later, these parameters were chosen because they were capable of capturing the amount of pure selectivity in the model (any lower value of the threshold would lead to too few cells with pure selectivity, for example). On the left are the percentage of cells with different selectivities as in Figure 3C. The bars are the data and the lines are the model. On the right, are histograms of model values from 100 instantiations, with the red markers showing the data values. The model matches the average firing rate and *FF*_*T*_ of the model, as it was fit to do so. Clustering, RV, and the amount of mixed selectivity are too low in the model. We use these parameters as the starting point for learning in this model.

## 3.3 Circuit Model with Hebbian Learning Captures PFC Responses

As described above, responses of PFC cells have a set of qualities that cannot be explained by random connectivity. In particular, the inability of the random network to simultaneously capture the values of response variability, clustering, pure, and mixed selectivity shows that PFC cells have a balance of specialization that may require learning to achieve. Here, we tested two variants of Hebbian learning to determine if a network endowed with synaptic plasticity can capture the elements of the data that the random network could not. The simple form of Hebbian learning that we use is based on the idea that the input populations that randomly start out giving strong inputs to a cell would likely make that cell fire and thus have their weights increased.

In both variants of learning tested, each cell has the weights from a subset (*N*_*L*_) of its input populations increased while the rest are decreased to keep overall input constant (this is done via a weight increase step and a normalization step). Such balancing of Hebbian and homeostatic plasticity has been observed experimentally (Keck et al., 2017), particularly via the type of synaptic up and down regulation used here (Chistiakova and Volgushev, 2009; Bourne and Harris, 2011; Scanziani et al., 1996; Lo and Poo, 1991). Therefore, it is plausible for an individual neuron to be able to implement such changes across its synapses.

The difference between our two variants of learning comes from which input populations are increased. In general, the top *N*_*L*_ input populations from which the cell already receives the most input have their weights increased (to capture the “rich get richer” nature of Hebbian learning). In the “constrained” variant, however, weight increases onto a PFC cell are restricted to populations of input cells that come from different task variables (e.g., cue 1 and cue 2. For a detailed explanation see Methods). This was done to ensure that cells had enough variety of inputs to create mixed selectivity. In the free variant, the populations from which a cell receives increased input due to learning are unrestricted. That is, they are determined only by the amount of input that the cell originally received from each population as a result of the random connectivity. This unrestricted form of learning is more biologically plausible as it can be implemented in a way that is local to the post-synaptic neuron, without knowledge of the identity of the upstream inputs. A toy example of each variant can be found in Figure 5B. In this example, free and constrained learning select different input populations to be enhanced, however, given random weights, free and constrained learning will select the same input populations in some cells.

Figure 5C shows how the weight matrix changes with different *N*_*L*_ values (the number of populations from which weights are increased during learning). Eventually, the learning leads to a steady state in which each PFC cell receives input only from cells in the top *N*_*L*_ populations. The higher the *N*_*L*_ the faster the matrix converges to its final state. When *N*_*L*_ is low, convergence takes longer as all the weight is transferred to a small number of cells. This plot is shown with a learning rate of.2.

The results of both forms of learning are shown in Figure 6A. The effects of learning are dependent on *N*_*L*_, and different *N*_*L*_ values are in different colors (*N*_*L*_ = 1, 2, 3 are tested here). Free learning is shown with solid lines, and constrained with dotted lines, except for the case of *N*_*L*_ = 1, where free and constrained learning do not differ and only one line is shown. In each plot, the data value is shown as a small black dotted line.

Clustering, mixed selectivity, and RV all increase with learning, for any value of *N*_*L*_ and both learning variants. When *N*_*L*_ = 1 (green line), mixed selectivity peaks and then plateaus at a lower value (as connections to all but one population are pruned), while other values of *N*_*L*_ plateau at their highest values. As it was designed to do so, constrained learning is very effective at increasing mixed selectivity, eventually getting to nearly 100 percent of cells. Free learning produces more modest increases in mixed selectivity, with *N*_*L*_ = 2 leading to slightly larger increases than *N*_*L*_ = 3. Before learning, the model matches the data’s balance of supra- and sublinear interaction effects (ratio of positive to negative terms: 1.100 *±*.048), and learning does not impact this balance (1.095 *±*.053, STDs over 20 random instantiations).

A factor impacting selectivity in this model—and especially with this task structure— is that cells that receive inputs from multiple populations from a single task variable may not end up having significant selectivity to that variable. This is especially true for the ‘task type’ variable, as cells can easily end up with input from both ‘recall’ and ‘recognition’ populations. If the inputs from these populations are somewhat similar in strength, the cell does not respond preferentially to either. This can help understand the discrepancy in how pure selectivity changes with free and constrained learning. In constrained learning, pure selectivity necessarily increases with learning (to the point where nearly all networks have 100% pure selectivity), whereas free learning can have inputs that effectively cancel each other out. A more direct investigation of how selectivity and other properties change with learning comes with the analysis of our toy model in the next two sections.

In these plots, both noise parameters are fixed, which allows us to see how *FF*_*T*_ varies with learning (this is also why the values at step 0 in Figure 6A do not always match those shown in Figure 4, as that model has noise parameters fit to match the data). The changes in *FF*_*T*_ stem from both changes in robustness to the additive noise and from changes in the mean responses, which impacts *FF*_*T*_ via the multiplicative noise term. Figure 6A shows that the variant of learning has less of an impact on *FF*_*T*_ than *N*_*L*_ does. In all cases, however, learning ultimately leads to lower trial variability in the model. This is consistent with observation made in PFC during training (Qi and Constantinidis, 2012).

Overall, low *N*_*L*_ leads to more acutely distributed weights and stronger structure and selectivity in the model. Constrained learning, with its guarantee of enhancing weights from different task variables, is also more efficient at enhancing structure and selectivity. The prefrontal cortex data shows a moderate level of structure and selectivity, therefore the approach that is best able to capture it is free learning with *N*_*L*_ = 3. In Figure 6B, we show how all of the model values compare to the data as this form of learning progresses. These plots, similar to Figure 4B and C, show values in units of standard deviations away from the model. It is clear from these plots that this form of learning leads all values in the model closer to those of the data. The best fit to the data comes after 6 learning steps with a learning rate of .2 (marked with a black arrow). At this point the ratio of the standard deviation to the mean of the weight distribution has only slightly increased, remaining within a biologically plausible range. While the best fit to the data comes before the model reaches its steady state, all values still eventually plateau to within *±*2.5 model standard deviations of the data. Furthermore, there are many reasons why PFC may not reach steady state; for example, once the animal’s performance plateaus, learning may slow (Glimcher, 2011). Also, other uses of PFC may interfere with learning and prevent the circuit from overfitting to this particular task. A detailed exploration of these mechanisms is beyond the scope of this study.

We plot the values of the data in comparison to the best-fit model in Figure 6C, similarly to Figure 4D. At this point, the average percent of cells with only pure selectivity is 25.4 *±*4.2, with only mixed 4.4 *±* 2.2, and with no selectivity 15.9*±* 4.1 (the comparable data values are ≈36%, 1%, and 13%, respectively). Thus, the model with learning is a much better fit to the data than the purely random network.

In addition to matching the measured properties of the PFC representation, we also tested if learning makes the neural representation more conducive to decoding. To do this for task information, we trained linear classifiers to readout out the task inputs (i.e., the identities of task type, cue 1, and cue 2 separately) as well as higher order terms (i.e., the combined identities of task type-cue 1, task type-cue 2, cue 1-cue 2, and task type-cue 1-cue 2). As expected from a higher dimensional representation, decoding performance is better in the population after learning, for both linear and higher order terms (Figure 6D, left). Post-learning accuracy for linear terms is 83.2% and for the higher-order terms 70.5% (the respective values for constrained learning after the same number of steps are 88.2% and 83%, not shown). We also used a same-different task to demonstrate how the representation after learning allows for better performance on a non-linearly separable problem. Here, all combinations of cue 1 and cue 2 identities were generated as inputs, and a linear classifier was trained to readout if the identities of the two cues were the same or not. Trying to read this information out from the input population is not very successful as these cells only have pure selectivity (Figure 6D, right). Random connectivity is sufficient to expand the dimensionality of the neural representations and to solve non-linearly separable problems. However, the model PFC population generated from random connectivity performs poorly because the low threshold that we determined by fitting the model to the data leads to low levels of mixed selectivity. After learning, the PFC population performs substantially better on this task.

## 3.4 Understanding Properties of Selectivity Before Learning

We have shown that Hebbian learning can impact selectivity properties in a model of PFC. Some of these impacts, particularly the increase in mixed selectivity, may seem counterintuitive. Here we use a further simplified toy neuron model to understand the properties of the network before learning and then demonstrate how learning causes these changes.

A schematic of this toy model is in Figure 7A, and it is described in the Methods. Briefly, the cell gets four total inputs–two (A and B) from each of two task variables (T1 and T2). The output of the cell is binary: if the weighted sum of the inputs is above the threshold, Θ, the cell is active and otherwise it is not. As in the full model, Θ is defined as a fraction, *λ*, of the sum of the input weights.

This format makes it easy to spot nonlinear mixed selectivity: if the cell is active (or inactive) for exactly one of the four conditions, it has nonlinear mixed selectivity to the combination of T1-T2. If the cell’s output can be determined by the identity of only one task variable, it has pure selectivity (and would be active for two of the four conditions). Otherwise it has no selectivity (active or inactive for all conditions) (see examples in Figure 2A and B).

Learning impacts selectivity by altering the way a cell represents these four conditions. To say more about how this occurs, we must first describe the properties of the representation in the random network before learning.

To be robust to noise, the cell’s response should be constant across trials within a condition. Additive noise can be thought of as a shift in the threshold, which may lead to a change in the cell’s response. Thus, trialwise additive noise drawn from a distribution centered on zero can be thought of as a range of effective thresholds centered on the original one (black dotted line in Figure 8A is the threshold without noise and the gray shaded area is the range of effective thresholds due to noise). If the inputs for a given condition fall in this range, the response of the cell will be noisy, i.e. flipping from trial to trial, and selectivity will be lost because the cell’s activity will not be a reliable indicator of the condition. Robustness to noise, then, can be measured as the range of thresholds a representation can sustain without any responses flipped, with a larger range implying higher noise robustness (if noise is drawn from a Gaussian distribution the noise range can represent thresholds within two standard deviations, for example, implying that a cell is robust to noise as long as its response is consistent on 95% of trials).

Assuming optimal threshold values (i.e., those with highest noise robustness) for each type of selectivity, the relative noise robustness of mixed and pure selectivity can be calculated (see Methods). We find that, thinking of the four conditions as the corners of a rectangle (as visualized in Figure 2C), mixed selectivity robustness depends on the length of the shorter side, while pure selectivity noise robustness depends on the difference between the two side lengths. We also find that, with random weights, most cells will have a representation that has higher noise robustness for pure selectivity than for mixed (see Methods).

Noise robustness changes, however, as thresholds deviate from optimal. The type of selectivity cells have in the absence of noise also varies with threshold in a related way. For example, using a low threshold may result in more cells with mixed selectivity and/or cells with pure selectivity that have low noise robustness (see Figure 2D for examples). To quantify these trends, we varied the threshold parameter *λ* and determined both the probability of different types of selectivity as well as the noise robustness for each type (see Methods for details). In Figure 7B, we show the fraction of cells that lose selectivity at a given noise level, for three different values of *λ*. Noise robustness (plotted as a function of *λ* in Figure 7C) is defined then as a normalized measure of the noise value that causes 50% of cells to lose selectivity.

Figure 7C demonstrates why the random network from which we start learning is necessarily in a condition of low mixed selectivity. Specifically, the value of *λ* we choose to use is constrained by the fact that the data shows high levels of pure selectivity. Therefore, we need a value that has high probability of pure selectivity and high noise robustness for it (especially because, as we will show, pure selectivity is unlikely to increase much with learning). Values of *λ* that meet this condition are not favorable for mixed selectivity. Therefore, the best we can do is choose a value of, for example, .4, where probabilities of pure and mixed selectivity are even, but pure has higher noise robustness (therefore effective rates of pure selectivity are higher). The fact that mixed selectivity is less noise robust than pure in the full model can be seen in Figure 4A.

Note that while the *λ* used for the random version of the full model shown in Figure 4D was around .27, that value is not directly comparable to the *λ* values in these plots for many reasons. First, the full model has 3 task variables, compared to the 2 used in the toy model. This means that, from the perspective of mixed selectivity for 2 ask variables, a given *λ* value will create a higher Θ in the full model with 3 task variables than in the toy one that has only 2 (because Θ is a function of the sum total of all weights, not just those relevant for the 2-way selectivity). In addition, in the toy model, 50% of the inputs are on for any given condition, whereas the nature of the task in the full model means that only 25% of inputs are on when looking at C1xC2 mixed selectivity, while one-third are on for TTxC1, TTxC2, and TTxC1xC2 mixed selectivity. The percentage of cells are also not directly comparable, as cells in the full model are labeled as pure if they have any of 3 different types of pure selectivity, and mixed if they have any of 4 different types of mixed. This toy model is thus meant to provide intuition only.

## 3.5 How Learning Impacts Selectivity

For the reasons just discussed, the random model starts in a regime where pure selectivity has high noise robustness and mixed does not. In order to match the amount of mixed selectivity seen in the data, we must then rely on learning to increase noise robustness for mixed selectivity, allowing more mixed cells to move out of the noise range.

Learning impacts noise robustness by expanding the representation of the different conditions. An example of this is in Figure 8A, where the gray shaded area represents the noise-induced range of the threshold. Before learning, the cell’s response is impacted by the noise. With learning, different conditions get pulled away from each other and the threshold, creating a much more favorable condition for mixed selectivity to be robust to noise. As can be seen, the responses are now outside the noise range. For the same reason that learning increases noise robustness (because the expansion increases the range of thresholds that support mixed selectivity), it can also increase the probability of a cell having mixed selectivity in the absence of noise. This can be seen in Figure 8C (left), where learning steps are indicated by increasing color brightness (constrained learning with rate of .25). At lower *λ* values, cells that are initially above threshold for all conditions (no selectivity) gain mixed selectivity with learning. But for *λ* values that support higher levels of pure selectivity (e.g., *λ* = .4, marked with a black dotted line), the percent of cells with mixed is not as impacted by learning. The percent of cells with pure selectivity increases only slightly at most *λ* values.

Noise robustness has a different pattern of changes with learning (Figure 8C, right). In particular, at *λ* = .4, the noise robustness still increases with learning even when the percent of cells with mixed selectivity doesn’t change. Furthermore, when starting from a *λ* value that has unequal noise robustness for pure and mixed selectivities, if most cells with pure selectivity are already robust to a given noise value, an increase in noise robustness for pure would only have a moderate effect on the population levels of pure selectivity. Conversely, if most mixed cells have noise robustness less than the current noise value, an increase in that robustness could strongly impact the population. In the same vein, a decrease in robustness will impact the pure population more than the mixed. Thus, changes in noise robustness seem to play a large role in the increase in mixed selectivity observed in the full model.

In particular, constrained learning with *N*_*L*_ = 2 always increases the lengths of both sides of the rectangle (as one weight from each task variable increases and the other decreases). As mentioned above, noise robustness for mixed selectivity scales with the length of the shorter side and so it necessarily increases with learning in this condition. Under certain weight conditions, noise robustness will also increase for cells with pure selectivity (this can be seen in Figure 8C, see Methods for details).

If *N*_*L*_ = 1, only one side length will increase and the other decrease. If the shorter side decreases, mixed selectivity noise robustness decreases. If the shorter side increases, mixed noise robustness increases, up until the point at which side lengths are equal. At that point the shorter side is now the decreasing side and mixed noise robustness goes down. This trend is reflected in the shape of the mixed selectivity changes seen with *N*_*L*_ = 1 in Figure 6A (mixed selectivity increases then decreases).

When using free learning (with *N*_*L*_ = 2), a portion of the cells will by chance have the same changes as with constrained learning. The remaining cells cause the differences observed between the two versions of learning, and can be of two types. In the first type, the larger side length increases and the smaller shrinks, causing a decrease in mixed noise robustness. Free learning doesn’t achieve the same levels of mixed selectivity as constrained because these cells continue to be too noisy. In the other type, the shorter side increases and the larger decreases, reducing the difference between the two side lengths and thus reducing pure noise robustness. Free learning loses pure selectivity as these cells become too noisy (as seen in 6A). More detailed descriptions of changes with learning can be found in the Methods.

Inputs from additional task variables can be thought of as a source of noise as well. In Figure 8B, we add a third task variable to the toy model. Now, in the case of the T1B-T2A condition, the identity of T3 determines if the cell is active or not. From the perspective of T1-T2 mixed selectivity, this has the same impact as shifting the threshold, and thus creates noise. If both T3 inputs are weaker than the strongest two inputs from T1 and T2 (as they are here), they will decrease with learning. This means that not only do different T1-T2 conditions get pulled apart with learning, but the same T1-T2 conditions become closer. This reduces the impact of “noise” from other task variables, and explains why mixed selectivity increases more with *N*_*L*_ = 2 than with *N*_*L*_ = 3 (Figure 6A).

In sum, learning changes a cell’s representation of the task conditions. Depending on the threshold value, this can create changes in the probability of mixed and pure selectivity and the relative noise robustness for each. Here, in order to match the high levels of pure selectivity seen in the data, we use a threshold regime where mixed selectivity noise robustness increases with learning. This causes a gain in the number of cells with mixed selectivity, such that it reaches the level seen in the data.

## 3.6 How Learning Impacts Other Properties

The visualization of this toy model gives intuition for why other properties change with learning as well. RV, for example, increases with learning (Figure 6A). The expansion that comes with learning places different conditions at different distances from the threshold. With a sigmoidal nonlinearity, this would translate to more variance in the responses across conditions, increasing RV. Because constrained learning ensures the most expansion, it increases RV more. These increases depend on *N*_*L*_ because lower *N*_*L*_ allows for a more extreme skewing of weights, and thus a subset of conditions will be far above threshold while the rest are below (leading to a high RV). RV has a limit, however, because even with *N*_*L*_ = 1, the cell would still respond equally to a quarter of the conditions (assuming an input from a cue variable)

Clustering values are also impacted by how selectivity changes. Clustering in the data appears to be driven by task type selectivity (Figure 3C), and as task type preferences develop in the model the clustering value increases. Here, the relative sizes of the input populations play a role. Because the input populations that represent task type contain more cells (Figure 5A), these populations are more likely to be among the strongest inputs to a cell, and thus have their weights increased (Note that this bias in favor of task type could also arise from the fact that only two task types are possible, and thus these inputs are on twice as often as cue inputs. Such a mechanism cannot be implemented in this model, however, so we use uneven numbers of input cells). Therefore, task type selectivity becomes common and clusters form around the axis representing the first regressor (which captures task type preference). This effect is weaker with free learning because both task type populations may have their weights increased, which diminishes the strength of task type preference. Lower *N*_*L*_, which minimizes preferences to other task variable identities, allows these clusters to be tighter.

Finally, it is important to note that the strength of inputs shown in Figures 2 and 8 (the colored arrows) correspond to, in the full model, the summed input from all cells representing a given task variable identity (i.e.,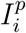), not just to weights from individual cells. These summed values are what need to change in order to expand the representation and see the observed changes. This is important for why the Hebbian procedure described here is effective at changing selectivity, as it assumes that many cells, acting in unison to cause post-synaptic activity, would lead to the increase of their individual synaptic weights, and thus an increase in the sum of those weights. Merely increasing the variance of the individual weights does not cause such a coordinated effect and would be less effective at driving these changes (as was shown in Figure 4C), especially with larger input population size.

## 4. Discussion

Here, motivated by several theoretical proposals about properties that would benefit encoding, we explored how prefrontal cortex represents task variables during a complex task. In particular we were interested in measures of selectivity (particularly nonlinear mixed selectivity), response density, and clustering of cell types according to preferences. By quantifying and measuring these properties in a PFC dataset, this work connects theoretical literature with experimental data to give insight into how PFC is able to support complex and flexible behavior. Furthermore, we explored how these response properties could be generated by a simple network model. Through this, we find evidence that the particular level of specialization and structure in the PFC response is not readily achievable in a random network without Hebbian learning. After Hebbian learning, the model—despite its relative simplicity—is able to capture many response properties of PFC. The changes that come with learning act via an expansion of the way cells represent conditions, and corresponding changes in noise robustness.

Interestingly, the variant of Hebbian learning that best matches the data is not the most effective at increasing mixed selectivity. It may be that the more effective method (”constrained” learning) would be too difficult to implement biologically, but perhaps there is also a computational benefit to the balance of mixed and pure selectivity found in the data. Particularly, preventing high levels of selectivity to this particular task may allow the network to retain flexibility.

In addition to retrospectively matching experimental results, this model also makes predictions regarding how certain values should change with training. In particular, clusters of cells defined by selectivity are expected to emerge with training and cell responses should become less dense across conditions. Previous work (Rigotti et al., 2013) has shown the value of mixed selectivity for the ability of a population to perform complex tasks. This work shows that mixed selectivity increases with learning, and these changes in PFC may correspond to increases in performance (Pasupathy and Miller, 2005), as learning in our model leads to increases in performance on classification tasks. Perhaps surprisingly, this model also predicts a concurrent, though small, decrease in pure selectivity. However, studies that have tracked PFC responses during training show signs of these changes. For example, in (Meyer et al., 2011), the amount of pure selectivity was measured directly pre- and post-training, and a significant drop in the percent of cells with pure selectivity was indeed observed. Furthermore, in hippocampus, an increase in mixed selectivity and slight decrease in pure was also observed with learning (Komorowski et al., 2009). In Meyers et al. (2012), the ability to readout match/nonmatch of two input stimuli from the population increases dramatically with learning, suggesting an increase in mixed selectivity. However, the ability to decode the identity of the stimuli (in the comparable portion of the trial) decreases slightly after training, which would be at odds with our linear classification results.

Our model makes many simplifying assumptions. The inputs, for instance, are binary cells that encode only the identity of different task variables. While this implies that the cells representing cue identities already have mixed selectivity (responding to the combination of the image and its place as either cue 1 or cue 2), it is still an assumption that the cells providing input to PFC are otherwise unmixed. This is something that, given current experimental evidence seems plausible (Pagan et al., 2013), but would benefit from further experimental exploration.

It may seem possible that adding more layers to the network would be a way to get the model to match the data without the need to introduce learning. This, however, is unlikely. For one, the data has high levels of pure selectivity which would be difficult to maintain through layers of random connections. Mixed selectivity, too, could decrease with layers, especially if each layer is noisy (which would be the realistic way to build such a model). It is also not obvious how such a model would achieve the clustering values observed in the data. Preliminary work on multi-layer models supports these intuitions (not shown). Also, such a model would not be able to address the changes with training discussed above. Finally, such a model would necessarily contain more parameters than a single layered network, and that would need to be taken into account when comparing to our learning model, which only introduces two additional parameters (*N*_*L*_ and the amount of learning, defined by the combination of learning rate and number of steps).

Another valuable endeavor would be to expand this model in the temporal domain. Currently in the model, all the task variable inputs are given to the network simultaneously. In the experiment, of course, there is a delay between cue 1 and cue 2. Delay activity is known to exist in areas like IT (Woloszyn and Sheinberg, 2009; Fuster and Jervey, 1982), and so this information could be being feed into PFC at the same time. But presumably, recurrent connections in PFC, and even possibly between PFC and its input areas, can enhance or alter selectivity. A recurrent model could also explore how PFC responses and representation vary over the time course of the trial, as recent experimental work has provided insight on this (Murray et al., 2016). Interestingly, recent work has demonstrated that Hebbian learning can be used to train recurrent neural networks on context dependent tasks (Miconi, 2017).

## 5. Acknowledgements

GWL was supported by a Google PhD Fellowship and NIH (T32 NS064929). SF was supported by the Gatsby Charitable Foundation, the Simons Foundation, the Kavli Foundation and the Grossman Foundation. EKM was supported by NIMH (NIMH R37MH087027) and MIT Picower Institute Innovation Fund. Thanks to Brian Lau for supplying code that implements the tests of uniformity on the hypersphere. The authors declare no competing financial interests.

